# Vegetation change and functional composition shifts in southwestern China during late MIS 3 to LGM

**DOI:** 10.64898/2026.06.25.734119

**Authors:** Kai Li, Ziru Hao, Pengyu Li, Xiao Zhang, Lina Liu, Mengna Liao, Zhenghong Tan, Yongbo Wang, Jian Ni

## Abstract

The climatic transition from Marine Isotope Stage 3 (MIS3) to the Last Glacial Maximum (LGM) had caused widespread vegetation change. Despite the dynamic equilibrium between vegetation and climate, the specific role of functional composition in vegetation’s response to climate change was inadequately understood. Here, we analyzed the long-term trajectories of palynological diversity, vegetation coverage and community-weighted-mean (CWM) functional traits based on EH22 pollen record (35–18 cal ka BP) from Erhai Lake, southwestern China. The results disclosed a vegetation transition from temperate deciduous broadleaf forest dominance in late MIS3 to cold coniferous and mixed broadleaved/coniferous forests in LGM. This vegetation dynamic involved functional composition shifts from competitive-driven functional convergence to partial recovery via niche differentiation during the late MIS3, and finally to a low-diversity but functional differentiation state through trait complementarity and diversification strategies during the cold LGM. Our results likely support a function-mediated climate filtering process whereby climate change regulated long-term vegetation dynamics during the MIS3–LGM transition primarily through shifts in CWM functional composition. These findings underscore the potential of pollen-based trait approaches to reconstruct ecosystem properties and advance our understanding of ecosystem change over decadal to millennial time-scales.

## Introduction

Glacial-interglacial cycles are the dominant theme of Quaternary, influencing the distribution, assemblage, and functional evolution of vegetation. Despite the unprecedented warming within current interglacial period, concerns about the rapid transition to glacial period exist (Wunderling et al., 2024). Marine Isotope Stage 3 (MIS3), as the most recent interglacial stage within the Last Glacial Cycle, features a climate similar to the present (Huber et al., 2006; Liang et al., 2022). The climatic transition from the late MIS3 to the Last Glacial Maximum (LGM) represents a process of cooling in the climate system, which may be one of the possible future scenarios faced by the current state of global warming. Global warming is significantly amplifying the global climate variability (Wu et al., 2025), and high climate variability is precisely one of the main characteristics of glacial climate (Rehfeld et al., 2018). Therefore, focusing on the vegetation change during the climatic transition from MIS3 to LGM will certainly benefit our understandings on the response of vegetation to high climate variability and provide a possibility of future ecosystem developments.

The MIS3 is featured by a warm climate but with significant fluctuations characterized by abrupt warming and extreme cooling (Cheng et al., 2016; Huber et al., 2006), leading to reorganization of regional vegetations. The early MIS3 fully developed forest remained a unique feature in central European pollen records, while with the cooling trend towards the glacial maxima of MIS2, tree pollen declined and steppe vegetation expanded (Kern et al., 2025; Sirocko et al., 2016). In China, the MIS3 vegetation likely bore a resemblance to that of today, but vegetation in eastern China had undergone alternating changes of forest and grassland between the wet and dry periods with climate fluctuations (Zhao et al., 2014). In contrast, MIS2 (approximately 27–11.9 ka BP) was generally cold and dry, in which Last Glacial Maximum (LGM) was the coldest period, with lowest atmospheric CO_2_ concentrations and the peak of global ice volume during the last Glacial Period (Clark et al., 2009). The harsh climate during LGM led to systematic adjustments in terrestrial vegetation. Global-scale vegetation reconstructions manifest that forests contracted significantly in most regions during the LGM, while the ranges of grasslands and shrublands expanded considerably (Li et al., 2024). The forests in northern China replaced by grasslands and desert steppes during dry periods, with frequent alternations in vegetation types in response to climate fluctuations (Ni et al., 2014). These widespread vegetation changes during the transition from MIS3 to LGM were likely caused by thermal deficiency, hydroclimate fluctuations, as well as habitat shrinkages. But the ecological mechanisms beneath the vegetation’s response to climate change remains less understanded.

Recently, pollen-based functional trait analysis provides an approach to disclose how functional composition varied along with climate change and disturbances (Birks, 2020; Brown et al., 2023). Palynological study is the most important way for studying long-term vegetation changes (Birks, 2019), and plant functional traits determine plant establishment, growth, and survival, and thus their responses to environmental change (Violle et al., 2007). By combining pollen analysis with plant functional traits, it allows botanists and ecologists to disclose the driving processes of long-term vegetation dynamics from the functional and ecological mechanism levels (van Der Sande et al., 2023). Despite certain limitation exist in pollen-based trait ecology (Birks, 2020; Brussel and Brewer, 2021; Liao et al., 2025), this approach has been sufficiently applied in tracking ecological function in trait, and reliability in extending the pollen-plant functional trait linkage into deeper time (Brussel et al., 2018; van Der Sande et al., 2023; van der Sande et al., 2019). Especially, pollen-based trait approach can not only identify the influence of different environmental factors on community structure and functional composition, but also explain the evolution of interspecies interactions and ecological strategies, thereby providing more mechanism-based and ecologically explanatory evidence for ancient vegetation changes (van Der Sande et al., 2023).

The extreme physiographical heterogeneity of southwestern China give birth to anomaly high species diversity of temperate plants (Xing and Ree, 2017), and making this region a hotspot of plant diversity (Myers et al., 2000). During the MIS3 to MIS2 stages, proxy-based records in the southwest of China manifested significant cooling trend (Lu et al., 2024; Zhang et al., 2023; Zhao et al., 2021), as a result of which, the vegetation composition underwent notable changes (Chen et al., 2014; Liao, 2019; Tang, 1992; Zhang et al., 2020). However, how vegetation responds to this cooling trend during the climate transition remains to be further explored. Specifically, pollen analysis mostly focuses on the vegetation compositional change or aim to quantitative reconstruction of climate, which overlooked the ecological process beneath the vegetation’s response to climate. Our current knowledge regarding the functional composition dynamics during the transition of MIS3 to LGM is still scarce. Here, we conducted pollen analysis on EH22 core sediment (35–18 cal ka BP) from Erhai Lake, southwestern China (Supplementary). By using Landscape Reconstruction Algorithm (LRA, Sugita, 2007), Biomization (Prentice et al., 1996) and pollen-based functional trait approach, we aim to (1) quantitatively estimate regional vegetation compositional change, (2) reconstruct community-level functional composition shifts, and (3) disclose how the vegetation structure and functions respond to climatic transition from MIS3 to LGM.

## Materials and methods

### Field work and pollen analysis

In May 2022, a 15.4-m-long sediment core of EH22 was retrieved from the center of Erhai Lake at a water depth of 20 m, by using UWITEC platform system and piston corer (Fig. 1). The age-depth model has been established based on 14 AMS ^14^C measurements (Hao et al., 2026), by using Bayesian modeling package “rbacon”(Blaauw et al., 2017), with the IntCal20 radiocarbon calibration curve (Reimer et al., 2020). Pollen analysis was conducted on a total of 215 samples from the 852 cm to 1,542 cm of EH22. About 1–2 g of sediment was pretreated with HCl (10%) and KOH (10%) to dissolve calcareous minerals and humic components, respectively. Pollen grains were then concentrated by heavy liquid flotation with ZnBr_2_ (2.2-2.3 g/mL). A tablet containing 10,315 *Lycopodium* spores was added to each sample for calculating pollen concentrations. Pollen identification was carried out with optical microscopy (ZEISS Axio.Scope.A1.), with a minimum of 400 grains counted for each sample. Stratigraphically constrained cluster analysis was used on terrestrial pollen to determine the biostratigraphy zones (CONISS, Grimm, 1987).

**Figure 1.**
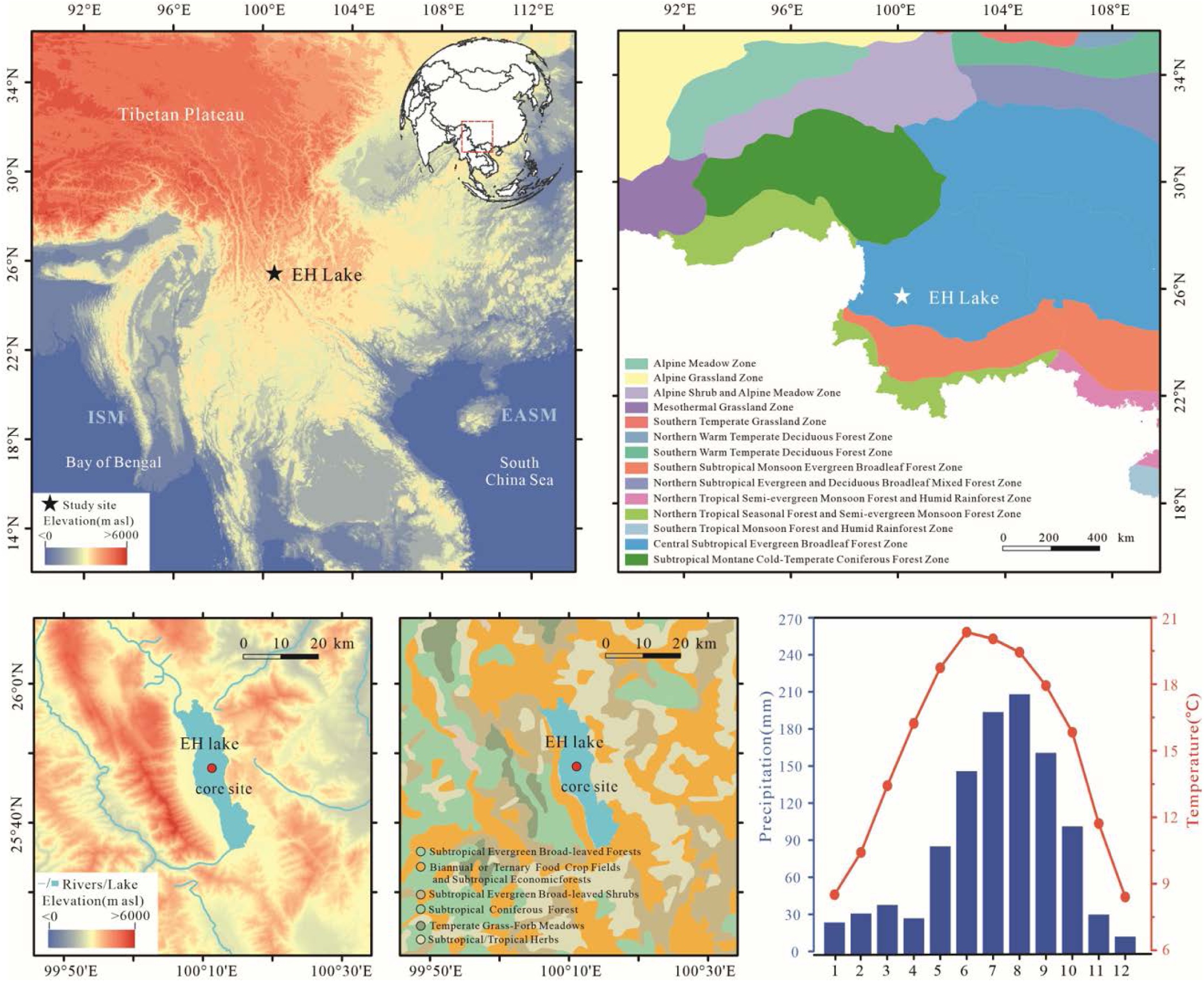
Map illustrating the location, topography, surrounding vegetation, and climatic characteristics of Erhai Lake. Vegetation maps from Zhang (2007).

### The REVEALS model and Biomisation

The REVEALS model was applied to quantitively reconstruct the relative vegetation coverage around Erhai Lake (Sugita, 2007). The existing relative pollen productivity (RPP) data from subtropical China and from the Northern Hemisphere were collected, with average value of all reliable RPP data for a certain type of pollen used as integrated RPP value. Poaceae was used as the reference taxon. Finally, the RPP values of 34 families and genera were ultimately selected in this study (Table S1). The cumulative percentage of these 34 taxa exceeded 90% in average in EH22. The fall speed of pollen (FSP) for every taxon was calculated according to Stoke and/or Falck formulas, based on pollen morphological characteristics in southwestern China (Tang et al., 2016). The calculations of pollen-based regional plant cover were conducted by using LRA.REVEALS.v6.2.2, with a constant wind speed of 3 m/s, and Z_max_ (maximum extent of the regional vegetation) of 100 km. The Principal Component Analysis (PCA) was then applied to explore the vegetation compositional change over time, by using R package *vegan* (Oksanen et al., 2001).

Biomization converts pollen information into biological groups of significant ecological importance, thereby inferential the vegetation types. Biomization was conducted with previously reported plant functional types (PFTs) (Ni et al., 2014) and the biome classifications for southern China (Zheng et al., 2023). Biomes include alpine shrub-land, meadow/tundra (ALSM), cold mixed forest (CLMX), temperate deciduous broadleaved forest (DBLF), subtropical evergreen broadleaved forest (EBLF), and tropical rainforest (TRFO). Modifications include using plant coverages as input data, adding a biome of evergreen sclerophyll forest (ESQF) representing the PFTs of evergreen sclerophyll *Quercus* (es) and a biome of hemlock forest (TSUF) comprising a PFTs of temperate conifer *Tsuga* (tsu). The detail assignment of pollen to each PFT and biome are given in Tables S2 and S3.

### Diversity measures and functional composition

To provide a mathematically rigorous basis for estimating richness and diversity of pollen, a series of diversity numbers following Hill (1973) were computed (Birks et al., 2016). Rarefaction of *Hill* numbers was carried out using the R package *iNEXT* (Hsieh et al., 2015), with a rarefaction count of 400 grains. The results included *N0* (richness), *N1* (exponential Shannon diversity or entropy measure), and *N2* (reciprocal Simpson’s diversity or concentration measure). To estimate temporal *β* diversity of pollen assemblage, we computed the Sørensen-dissimilarity value (the total dissimilarity, *β*_sor_), the nestedness fraction of the Sørensen-dissimilarity value (nestedness component, *β*_sne_), and the Simpson-dissimilarity value (turnover component, *β*_sim_) for the adjacent pairs of samples in each record. *β*_sor_, *β*_sne_ and *β*_sim_ were calculated using R package *betapart* (Baselga et al., 2012). In order to highlight the main trends, the LOESS smooth method was applied on *Hill* numbers and *β* diversity.

Plant trait data of modern vegetation were obtained from published Chinese plant trait datasets (An et al., 2024; Wang et al., 2022), and our own fieldwork on Hengduan Mountains and Tibetan Plateau (Jin et al., 2023). Only data from southwestern China were selected in consideration of accuracy of habitat and function (Supplementary Figure S1). Plant height, seed mass (SM), leaf thickness (LT), leaf area (LA), specific leaf area (SLA), leaf mass per area (LMA), leaf dry matter content (LDMC), leaf carbon content (LCC), leaf nitrogen content (LNC), and leaf phosphorus content (LPC) were selected. These traits are with ecological significance such as resource acquisition, environmental adaptation and reproductive strategies (Supplementary Table S4). We initially calculated the mean trait value for each plant family and genus, and the results were assigned to the corresponding pollen taxa in each fossil sample. Community weighted means (CWM) were used to reconstruct past vegetation trait changes, weighted by plant coverage that computed from pollen percentage data with LRA calibration (REVEALS here). Functional dispersion (FDis) was computed as an indicator of the functional diversity exhibited by species across a multidimensional trait space (Laliberté and Legendre, 2010). High FDis values indicate a greater distribution and functional differentiation of species, in terms of their functional characteristics, implying a more diverse range of functions within ecosystems. Ultimately, a LOESS method was utilized to smooth changes of CWM traits and FDis over time. The calculations were carried out using the R package *FD* (Laliberté et al., 2009).

### Statistical analysis

To disclose the relationships between vegetation characteristics (richness, FDis, PCA_REVEALS_ and *β*_sor_) and climate forcings, mental test was firstly applied. Climate variables include global atmospheric CO_2_ concentration (Bereiter et al., 2015), total solar irradiation (TSI) on July and January at 25°N (Laskar et al., 2004), sea surface temperatures (SST) from tropical Indican Ocean (SO189-119KL, Mohtadi et al., 2014), Asian Summer Monsoon (ASM) intensity of stalagmite δ^18^O from Chinese caves (Cheng et al., 2016), and brGDGTs-based mean annual temperature (MAT) from Tengchong Qinghai Lake (Zhao et al., 2021). Mantel test was conducted by using the *mantel_test()* function from the R package *linkET* (Huang, 2021).

Structural Equation Model (SEM) based on a maximum likelihood method was then introduced to explore the interrelationships among climate change, functional composition shifts, and vegetation change (PCA_REVEAL_). Climatic variables with |*r*| > 0.80 were removed to eliminate strong collinearity coefficients, yielding ASM (stalagmite δ^18^O), atmospheric CO_2_ and MAT as climate forcings. PCA analysis was conducted on all CWM traits to extract main functional compositions and reduce their collinearity. Based on the PCA loading matrix and their ecological significance (Table S4; Fig. S2), CWMs for LA, LMA and LCC were selected to represent the main functional dimensions of vegetation. A network structure (climate-function, climate-vegetation, function-vegetation) was first reconstructed without defining structural paths among latent variables. We refined the model structure by adding potentially significant pathway(s) and/or removing those with extremely weak standardized path coefficient to improve the fit. The minimal and best model was selected based on the Akaike Information Criterion (*AIC*). *Fisher’C* test was conducted to assess the global goodness of model fit, with *P* ≥ 0.05 indicating reliable fits. SEM analysis was conducted using the R package *piecewiseSEM* (Lefcheck et al., 2015). Standardized path coefficients were used to measure the direct and indirect effects of predictors. Direct effects were measured as the standardized path coefficients between variables, while indirect effects were calculated as the product of direct path coefficients along a specific causal chain.

## Results

### Pollen compositional changes

A total of 80 pollen taxa were identified, including 49 families and 31 genera. Arboreal pollen was dominant. The coniferous taxa mainly comprise *Pinus*, *Picea*, *Abies*, *Tsuga*, and Cupressaceae. Evergreen broad-leaved taxa mainly include evergreen *Quercus* and *Castanopisis*. Deciduous broad-leaved taxa were dominant by deciduous *Quercus*, *Alnus*, *Betula*, *Corylus*, *Ulmus*, *Carya* and *Juglans*. The shrub taxa mainly consist of Rosaceae, Ericaceae, and Oleaceae. Herbaceous pollen taxa were abundant, comprising mainly Poaceae, *Artemisia*, Cruciferae, Lamiaceae, and Asteraceae. Six zones had been identified according to the CONISS result (Fig. 2):

**Figure 2.**
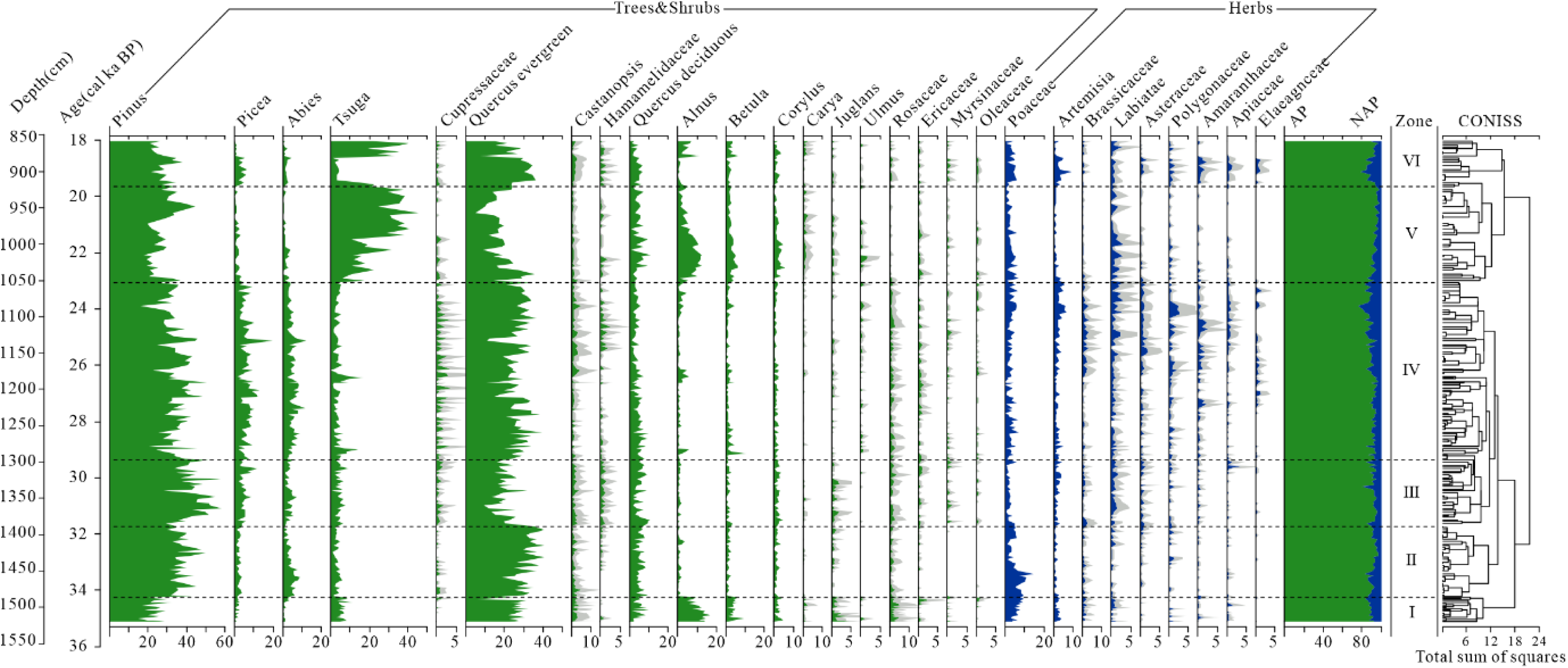
Pollen diagram showing the main taxa in EH22. The solid and faded plots show the same percentage values on actual percentages and × 3 exaggeration. AP, arboreal taxa pollen; NAP, non-arboreal taxa pollen.

Zone I (15.42–14.77 m, 35.2–34.3 cal ka BP). *Pinus* (23.7%) was dominant, and percentages of *Picea* (1.25%) and *Abies* (1.13%) were low. The evergreen *Quercus* (25%) and *Alnus* (11.2%) appeared as relatively high proportions. *Tsuga* (6.15%), deciduous *Quercus* (4.61%), and *Betula* (3.95%) were abundant. Shrub taxa mainly comprised Rosaceae (2.78%). Poaceae (6.08%) and *Artemisia* (2.36%) were abundant.

Zone II (14.77–13.58 m, 34.3–32 cal ka BP). *Pinus* (37.10%) increased, as well as *Picea* and *Abies*, whiles the content of *Tsuga* (3.14%) decreased. Evergreen *Quercus* increased to 31.8%. While most of the deciduous pollen remained stable, the percentage of *Alnus* decreased obviously. Shrub taxa decreased compared with previous zone. Poaceae (5.66%) still dominated the herbaceous pollen, with slightly increased Asteraceae.

Zone III (13.58–12.51 m, 32–29.4 cal ka BP). *Pinus* (43.36%) peaked, and *Picea* (4.98%) and *Tsuga* (4.35%) increased as well. The evergreen *Quercus* (20%) decreased obviously. Shrub taxa of Rosaceae (1.42%), Rhododenaceae, and Oleaceae slightly increased. Poaceae decreased but still dominated the herbaceous taxa.

Zone IV (12.51–10.04 m, 29.4–23 cal ka BP). *Pinus* (34.63%) and *Tsuga* (3.96%) decreased. *Picea*, *Abies* and Cupressaceae increased. Evergreen *Quercus* (26.57%) increased, as well as Hamamelidaceae, *Alnus*, and *Betula*. The relative abundance of *Castanopsis*, deciduous *Quercus,* and Rosaceae slightly decreased. Nearly all the percentages of herbaceous taxa increased in this zone.

Zone V (10.04–9.10 m, 23–20 cal ka BP). *Pinus* decreased to 27.16%, while *Tsuga* sharply increased to an average of 24.05%. *Picea* (1.54%) and *Abies* (1.49%) decreased. Evergreen *Quercus* (14.7%) decreased, while the deciduous taxa of *Alnus* (6.35%), *Betula* and Corylus increased. Shrub and herbaceous taxa decreased, with Poaceae and Labiatae dominant in herbaceous taxa.

Zone VI (9.10–8.52 m, 20–18 cal ka BP). *Pinus* decreased, as well as *Tsuga*, whiles *Picea* and *Abies* slightly increased. Evergreen *Quercus* (25.93%) and *Castanopsis* (1.4%) increased obviously. Nearly all the herbaceous taxa increased.

### Vegetation coverage and biome changes

Both of REVEALS-based coverage and biome changes illustrated clear vegetation change trajectory during the climate transition from late MIS3 to LGM (Fig. 3). Before 32 cal ka BP, the coverage of coniferous forests expanded in lowland region (pine forest) and high altitude (*Abies*/*Picea*), along with high coverage of sclerophyll oak forest and grassland (Poaceae). Biomes of ALSM, CLMX and DBLF appeared as high scores, indicating the dominance of alpine vegetations. During 32–27 cal ka BP, the coverage of alpine coniferous forest (*Abies*/*Picea*, and Cupressaceae) expanded obviously. While the CLMX remained domination, during 27–23 cal ka BP, the coverage of alpine coniferous forest shrank gradually. After 23 cal ka BP, the DBLF increased obviously, mainly caused by the coverage expansions of *Alnus* and deciduous oaks. Of note, the dominance of DBLF was gradually replaced by TSUF along with the expanding coverage of hemlock forest. A Rapid vegetation reorganizations occured since 21 cal ka BP, manifested by rapid replacement between TSUF, CLMX, and DBLF (Fig. 3). The principal variation of vegetation coverage manifested the gradual expansion of coniferous forests and shrinkage of broadleaved forests, as suggested by PCA result (Fig. S3). When approaching to the end of LGM, rapid vegetation reorganization between alpine coniferous forest (*Abies* and *Picea*) and temperate deciduous and coniferous forests (*Tsuga* and *Alnus*) appeared.

**Figure 3.**
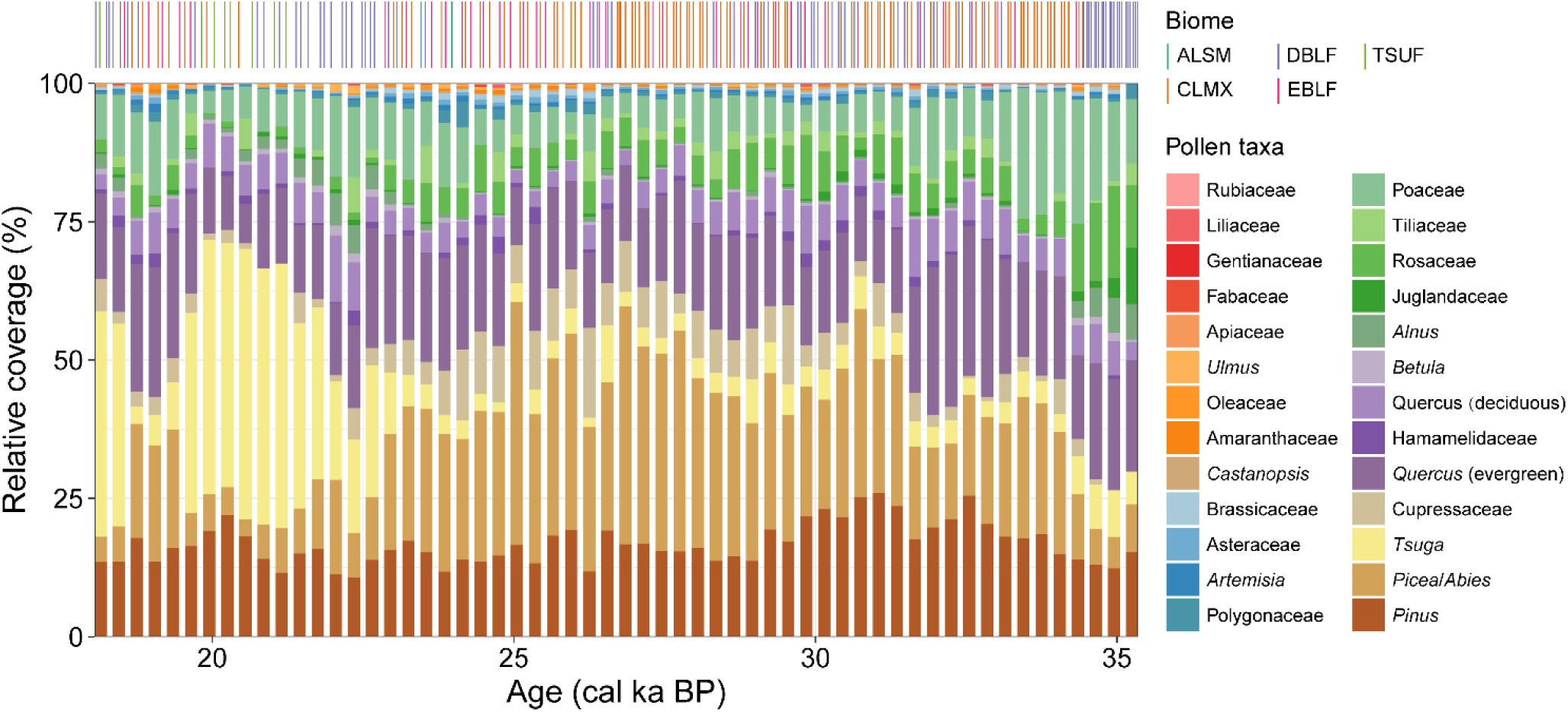
REVEALS model and Biomization outputs manifesting the changes in vegetation coverage and biome types in Erhai region during 35.2–18 cal ka BP. Best match biomes including ALSM, alpine shrub-land, meadow/tundra; CLMX, cold mixed forest; DBLF, temperate deciduous broadleaved forest; EBLF, subtropical evergreen broadleaved forest; TSUF, hemlock forest.

### Diversity and functional composition change

The palynological diversity in EH22 showed obvious fluctuations. The taxa richness (*N0*) was increasing during late MIS3, followed by a long-term decreasing trend within the LGM (Fig.4). The diversity indices of *N1* and *N2* decreased obviously before 32 cal ka BP, and after 23 cal ka BP. The total *β* diversity (*β_sor_*) showed long-term increasing trend since 35.2 cal ka BP, with exception during 30–25 cal ka BP (Fig. 4). The turnover component (*β_sim_*) increased before 30 cal ka BP and fluctuated after that. The nestedness component (*β_sne_*) progressively increased. The *β_sim_* was higher than *β_sne_*, manifesting that the taxa lost/gain was more important than that of taxa nesting. The *β_sne_* increased obviously after 24 cal ka BP and contributed equally with that of *β_sim_* at 18 cal ka BP, which suggested vegetation reorganization (compositional adjustment but not replacement) playing a crucial role in vegetation dynamics at the end of LGM.

**Figure 4.**
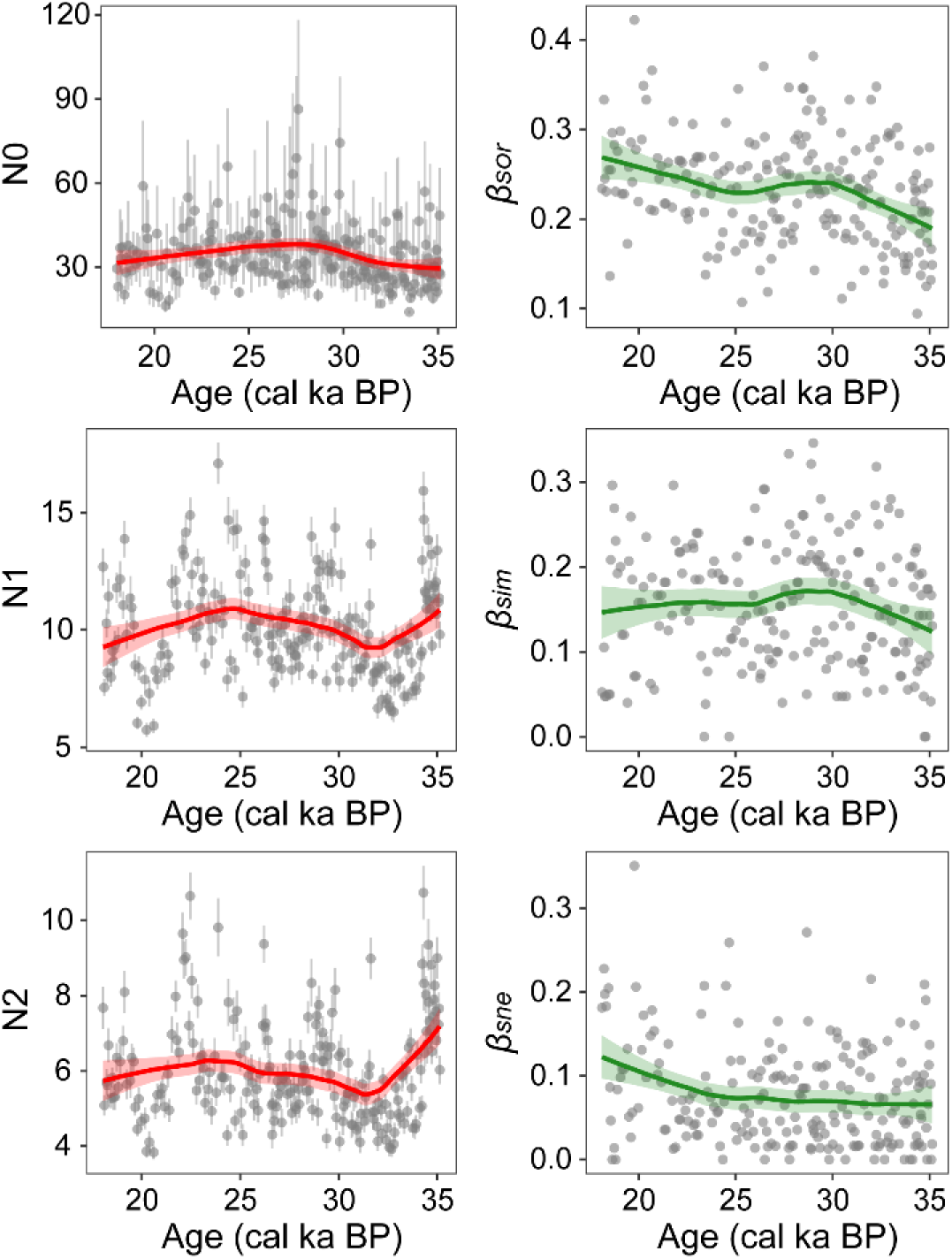
Palynological diversity of EH22, with species richness (*N0*), Shannon-Wiener (*N1*) and Simpson (*N2*) diversity indices, total beta diversity (*β_sor_*) and its additive partitioning into species turnover (*β_sim_*) and nestedness (*β_sne_*) components.

The CWMs of functional traits showed fluctuations during 35.2–18 cal ka BP, with obviously changes occurring at 27–25 cal ka BP (Fig. 5). The CWM of plant height (CWM_Height) and decreased slowly in the late MIS3 and increased rapidly in the LGM. The CWM of LA (CWM_LA) decreased rapidly before 32 cal ka BP, remained stable in 32–25 cal ka BP and followed by a decrease trend after that. The CWM of LMA (CWM_LMA) showed increasing trends in the late MIS3, contracting a rapid decreasing in the LGM. The CWM of LT (CWM_LT) was similar with LMA, but with slight decreasing in 32–25 cal ka BP. The CWMs of LDMC (CWM_LDMC) and LCC (CWM_LCC) declined before 32 cal ka BP, and increased after that. Long-term decreasing trends of CWMs of LNC (CWM_LNC), LPC (CWM_LPC), and seed mass (CWM_SM) had been disclosed, while the CWM_LPC manifested slight increasing trend within the LGM. Rapid increase of CWM of SLA (CWM_SLA) was clear after 25 cal ka BP. Before 32 cal ka BP, the FDis was high but with sharply decreasing, revealing strong ecosystem functional heterogeneity but with continued contraction of the community functional space (Fig. 5). After 32 cal ka BP, FDis exhibited a short-term upward trend and then a decline trend. After entering the LGM, the FDis decreased slightly, but turned to increase after 25 cal ka BP, indicating a contraction and then an expansion of functional compositions.

**Figure 5.**
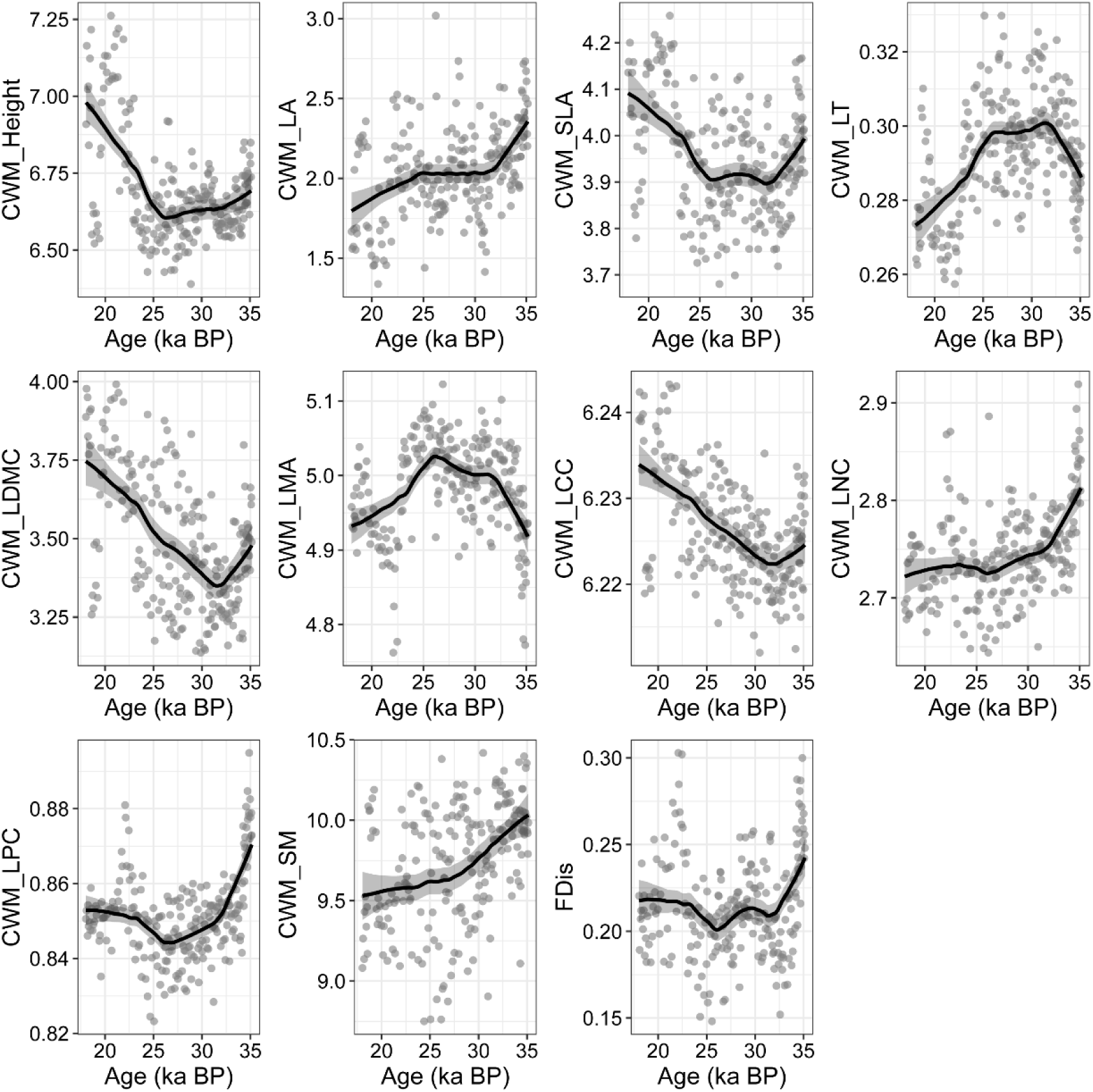
Community-weighted mean (CWM) of functional traits and FDis calculated from EH22, weighted by REVEALS-based vegetation cover.

### Relationships among climate dynamic, functional shift and vegetation change

Mantel analysis revealed that climate change was significantly correlated to vegetation change, *β_sor_*, and FDis (Fig. 6). Both of the PCA of REVEALs (REV_PC1) and FDis were significantly correlated with ASM, MAT, atmospheric CO_2_, and TSI on July and on January. There was no correlation between FDis and global mean temperature (NGRIP δ^18^O here). These findings indicated that local climate dynamics not only impacted the vegetation compositional changes but also affected the shifts of functional compositions. Meanwhile, there was no correlations between these climate parameters and palynological richness, and *β_sor_* was correlated only with atmospheric CO₂.

**Figure 6.**
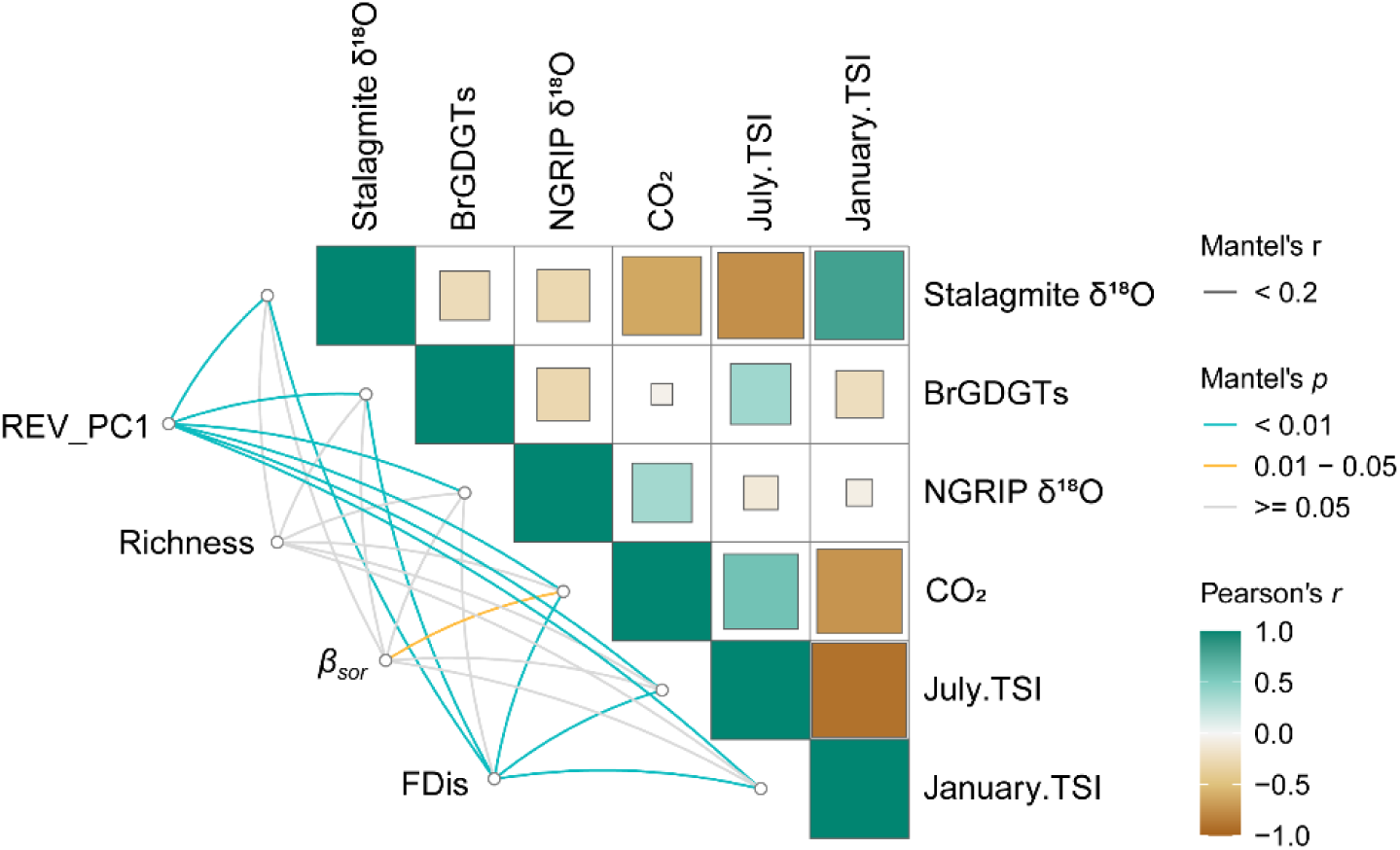
Mantel test between vegetation change and climate forcings of Asian Summer Monsoon Index (Stalagmite δ^18^O, Cheng et al., 2016), mean annual temperature from Tengchong Qinghai Lake (Zhao et al., 2021), Northern Hemisphere temperature (North Greenland Ice Core Project members, 2004), surface water temperature of SO189-119KL in the equatorial Indian Ocean (Mohtadi et al., 2014), total solar irradiation on July and January at 25°N (Laskar et al., 2004), and global atmospheric CO_2_ concentration (Bereiter et al., 2015)

The SEM performed well, with *Fisher’s C* of 15.412 (*P* = 0.351) and Chi-squared of 8.416 (*P* = 0.297). Standardized path coefficients revealed distinct associations among climate forcings, the CWM functions and vegetation change. The CWM_LCC (*R*^2^ = 0.17) was positively correlated with ASM (δ¹⁸O here, 0.281, *P* < 0.001), while negatively affected by MAT (BrGDGTs here, −0.239, *P* < 0.001). The CWM_LA (*R*^2^ = 0.13) showed a positive response to atmospheric CO_2_ (0.356, *P* < 0.001). The CWM_LMA (*R*^2^ = 0.3) had a positive correlation with MAT (0.421, *P* < 0.001) and a negative relationship with atmospheric CO_2_ (−0.328, *P* < 0.001). The total explanation of climatic forcings and functional shifts on vegetation compositional change (PC1 here) was 56%, in which the vegetation compositional change was positively affected by the atmospheric CO_2_ (0.378, *P* < 0.001) and community’s functional shifts of CWM_LCC (0.219, *P* < 0.001), and negatively impacted by CWM_LMA (-0.516, *P* < 0.001) and LA (-0.312, *P* < 0.001). Direct effects from CWM_LMA (-0.52), LA (0.312), and CWM _LCC (0.219) were clear. There was no direct effect from ASM and MAT to vegetation change, while their indirect effects were 0.062 and -0.27, respectively. The direct effect of atmospheric CO_2_ was 0.378 and its indirect effect was 0.059.

## Discussion

Our result illustrated obvious vegetation changes including palynological diversity, vegetation coverages and functional compositions at Erhai region, southwestern China, during 35.2–18 cal ka BP, which can be roughly divided into two stages: the late MIS3 (35.2–28 cal ka BP) and the LGM (28–18 cal ka BP). This phased difference is not only reflected in the fluctuation of plant composition, but also in the systematic transformation of community functional compositions, along with the long-term climate cooling in the southwestern China (Lu et al., 2024; Zhang et al., 2023; Zhao et al., 2021).

### Vegetation compositional change during the transition from MIS3 to LGM

EH22 pollen record manifested that the cold-adapted vegetation dominated during the climatic transition from MIS3 to LGM, with cold coniferous forests, temperate deciduous broad-leaved forests, and alpine meadow/shrub belts all maintaining relatively high proportions, while the proportion of subtropical vegetation types was gradually suppressed. During the late MIS3, the vegetation surrounding Erhai Lake was dominated by temperate deciduous broad-leaved forests represented by deciduous oaks, *Alnus*, and *Betula*, accompanied by herbaceous communities of Poaceae and *Artemisia*. As temperatures declined, temperate deciduous broad-leaved forests gradually diminished, herbaceous taxa increased, and cold coniferous vegetation dominated by *Picea*/*Abies* progressively gained dominance (Fig. 3a). Vegetation types transitioned from deciduous broad-leaved forests toward alpine coniferous forest that better suited to cold environments (Fig. 3b). This trend aligned with regional pollen records in southwestern China (Chen et al., 2014; Liao, 2019; Tang, 1992; Zhang et al., 2020). For instance, deciduous broad-leaved forests prevailing during the late MIS3 declined continuously as the climate gradually cooled, while that of evergreen sclerophyllous oak forests increased in Tengchong (Zhang et al., 2020). Similar pattern was also observed at the Lugu Lake and Xingyun Lake, where both cold coniferous forests dominated by *Picea*/*Abies* and cold- and drought-tolerant herbaceous groups dominated by Poaceae, *Artemisia*, and Brassicaceae expanded continuously since 29 cal ka BP (Chen et al., 2014; Liao, 2019). Collectively, these results indicate that vegetation changes across southwestern China during the transition from MIS3 to LGM followed a consistent pattern, characterized by a shift from temperate deciduous broad-leaved forests to cold coniferous forests and alpine meadow/shrubland.

Although the vegetation change around Erhai Lake resembled regional pollen records during the MIS3 to LGM transition, it exhibits a relatively unique feature at the late stage of LGM. It was outstanding that the hemlock forests expanded significantly and rapidly during 23–20 cal ka BP, reaching a peak at approximately 20.5 cal ka BP, while evergreen sclerophyllous oak forests and cold coniferous forests declined (Fig. 3a). Similar *Tsuga* expansions have also been recorded at Qilu Lake (Long et al., 1991) and Tiancai Lake (Xiao et al., 2014), although the changes were less pronounced than that around Erhai Lake. By using species distribution model (SDM) to study the distribution changes of *Tsuga* in Hengduan Mountains, we confirmed that regional climate change is able to cause a rapid reorganization of the vegetation in the late LGM (Liao et al., under review). Furthermore, the time points when the vegetation in Erhai Lake underwent rapid changes were in sync with Atlantic meridional overturning circulation (AMOC) anomalies, indicating that climate change is one of the primary factors driving the rapid changes in the vegetation.

### Community-level functional composition dynamics

Our result disclosed the functional composition shifts in response primarily to climate transitions from MIS3 to LGM. The diversity decline before 32 cal ka BP might have resulted from competitive exclusion and the formation of dominant taxa leading to functional convergence (Fig. 4 & 5). After 32 cal ka BP, diversity began to increase, probably suggesting community reorganization and structural complexification. The increased *β_sor_* and *β_sim_* likely reflected an overall increase in taxa turnovers (Baselga, 2010). Meanwhile, changes in FDis reflected a phased shift in the functional structure of community from rapid contraction to stabilization (Fig. 5). The decline of FDis before 32 cal ka BP indicated that environmental filtering or intensified competition led to functional convergence and a shrinking functional space; the subsequent stability indicated that communities re-expanded their functional space through trait differentiation and functional complementarity, enhancing resource utilization efficiency (Díaz et al., 2016; Laliberté and Legendre, 2010). These shifts implied that the dominant drivers of community assembly transitioned from competition and environmental filtering to functional complementarity and niche differentiation (Díaz et al., 2016). Changes in CWM traits further revealed adjustments in community functional compositions (Fig. 5). The continuous increases in LMA, coupled with the gradual decline in CWM_SLA and the trend of LDMC first decreasing then increasing, indicated a gradual shift from resource-acquisitive to conservative strategies (Gratani et al., 2018; Wilson et al., 1999). The synchronous declines in CWM LA, LNC and LPC suggests a weakening of photosynthetic and nutrient turnover rates, with plants increasingly relying on extended leaf lifespan and improved resource use efficiency to maintain functional stability (Reich, 2014; Wright et al., 2004). The increased LCC and LDMC after 32 cal ka BP, with lower SLA, LNC, indicated a resource-conservative growth strategy (Prieto et al., 2018). The SM declined, indicating that the community invested less on reproductive strategy under a steady-state structure (Mueller et al., 2024; Saatkamp et al., 2019).

Vegetation showed functional conservatism under the stress of a long-term cold condition during the LGM. The continuous decline in taxa richness and slight increases but continuous declines in *H1* and *H2* indicated that the vegetation gradually transitioned toward a low-diversity state dominated by a few regnant taxa (Fig. 4). It is likely that the community underwent taxon filtering and niche contraction, in which the temporal community differences intensified as suggested by the increased *β_sor_* during the LGM (Baselga, 2010). This temporal community difference primarily driven by the strengthening of nestedness structure (Fig. 4). The FDis decreased after entering the LGM, followed by slightly increasing trend (Fig. 5), indicating that, after filtering-induced convergence, the community’s functional differentiation restored partial stability and resource use efficiency through trait complementarity and diversification strategies (Adeleye et al., 2023; Laliberté and Legendre, 2010). The decreased LA, but continuously increased LCC and LDMC supported a resource-conservative growth strategy within cold LGM, under which plants tend to undergo more efficient carbon fixation with limited resource availability in order to reduce resource consumption (Han et al., 2024; Prieto et al., 2018). It was worth noting that the plant height showed obvious increase after 25 cal ka BP, suggesting a strengthen of community structure (Falster and Westoby, 2003). The high LMA and LT, with low SLA before 25 cal ka BP reflected the community’s maintenance of thick, dense leaf structures under strong environmental stress to enhance tolerance and resource conservation (Gratani et al., 2018; Wilson et al., 1999). As the increased plant height potentially received more sunlight and carbon dioxide, and consequently photo-synthesized more (Falster and Westoby, 2003), the community’s photosynthetic and nutrient acquisition capacity strengthened. This aligned well with the increases in SLA and LPC and decreases in LMA and LT (Fig. 5), exhibiting a brief resource-acquisitive strategy (Reich, 2014).

Given above, vegetation functional compositions shifted from competitive-driven functional convergence to partial recovery via niche differentiation during the late MIS3, with a gradual transition toward resource-conservative strategies responding to climate colling progress. In contrast, the vegetation transformed into a low-diversity but functional differentiation state through trait complementarity and diversification strategies during the cold LGM.

### Functional composition shifts mediated vegetation responding to climate change

The essence of vegetation change lies in the mortality and regenerate of individual plants, which drive alterations in community structure and composition, ultimately manifesting as temporal and spatial shifts in vegetation assemblages. Our findings demonstrate that, although different taxa and temporal vegetation were subjected to climate-change stress, it might be more important that the community adjusted their functional composition in harmony with long-term climate trends (Fig. 7). The late MIS3 was characterized by relatively warm and humid climate, while during the LGM, regional temperatures, humidity and atmospheric CO₂ concentration declined (Bereiter et al., 2015; Cheng et al., 2016; North Greenland Ice Core Project members, 2004). Accordingly, broadleaf taxa gained dominance during mild climatic phases through fast-economics functional traits, whereas coniferous taxa maintained long-term community stability during the cold LGM through conservative functional traits.

**Figure 7.**
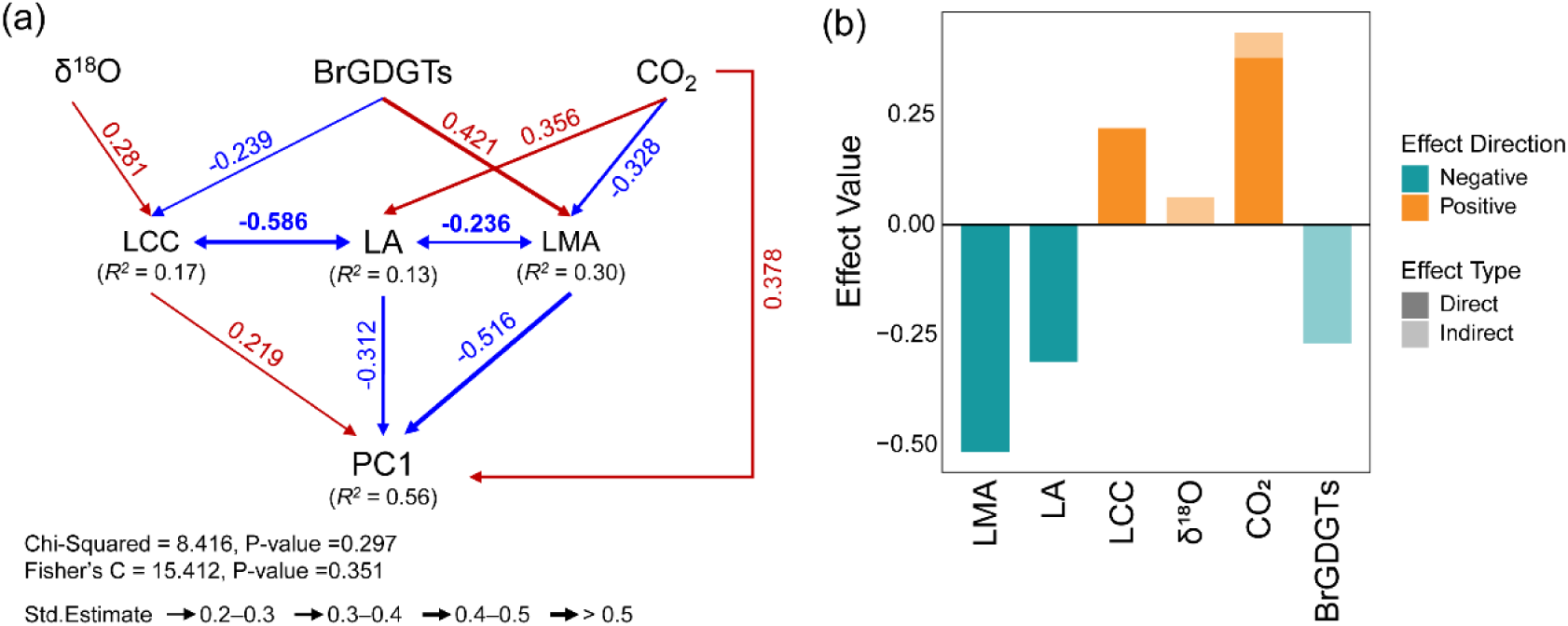
SEM analysis illustrating the significantly relationships (labeled coefficients) among climate factors, functional compositions, and PC1 of REVEALS-based vegetation change (a). The effects of CWM functional compositions (LCC, LA and LMA), atmospheric CO_2_ concentration (Bereiter et al., 2015), Asian Summer Monsoon Index (Stalagmite δ^18^O, Cheng et al., 2016), and mean annual temperature from Tengchong Qinghai Lake (BrGDGTs, Zhao et al., 2021) shown in (b).

Community-level functional composition dynamic reflects the overall functional characteristics of the community (Muscarella et al., 2017), serving as the integrated outcome of prolonged climatic and environmental changes (Shipley et al., 2006). The CWM functional composition characterizing the optimal functional strategy of plant communities is determined by the dominant species according to the mass-ratio hypothesis (Grime, 1998). Therefore, a few key species drive community adaptabilities under climate change (Dobson et al., 2026). Taxa of *Alnus* and *Betula* are featured by high LA but low LMA, indicating that when they dominated around 35 and 22 cal ka BP, the community tended toward lower per-unit leaf area construction cost—a typical fast resource-acquisition strategy (Reich, 2014; Wright et al., 2004). Meanwhile, they maintained a certain degree of leaf longevity and defense capacity through moderately high LDMC, consistent with the general pattern of fast-economics broadleaf trees dominating (Bruelheide et al., 2018; Díaz et al., 2016). In contrast, coniferous taxa tend toward a resource-conservative strategy. *Tsuga*, tall and with high LDMC, tends to develop a structurally stable, slowly regenerating canopy, aligning with that the hemlocks community grew tall to maximize light interception and enhance photosynthetic efficiency during the late LGM (Falster and Westoby, 2003; Wilson et al., 1999). *Picea*/*Abies* are with high LMA, manifesting that cold coniferous forest tended to enhance leaf resistance by increasing per-unit leaf area construction cost (Poorter et al., 2009; Villar et al., 2013). The expansion of *Picea*/*Abies* represented an extremely conservative leaf economic strategy of community adapted to glacial cold environments, helping to preserve high leaf physiological integrity and inter-annual survival under prolonged low-temperature stress. Thus, vegetation likely achieves climatic equilibrium by adjusting the functional composition of the community, a process manifested as shifts in species composition.

Our results likely support a function-mediated climate filtering process whereby climate change regulated long-term vegetation dynamics during the MIS3–LGM transition primarily through shifts in CWM functional composition. The SEM results show that precipitation and temperature factors affected vegetation change in the Erhai region mainly via their effects on functional composition (Fig. 7). This deduction is consistent with the well-established role of climate in regulating plant functional traits (Díaz et al., 2016; Li and Prentice, 2024). Moreover, pollen-based trait approaches are capable of capturing such CWM functional shifts along vegetation gradients (van der Sande et al., 2021), which lends objectivity to the climate–function correlations observed in our SEM. Climate change can drove vegetation compositional change by triggering plant mortality and recruitment. However, this process might operate indirectly: climate change reshapes the functional composition of the community—a process we term *functional filtering*—whereby only species with traits matching the prevailing environmental threshold maintain dominance. This mechanism steers species turnover and community assembly, ultimately manifesting as long-term changes in vegetation composition and structure. These findings underscore the potential of pollen-based trait approaches to reconstruct ecosystem properties and advance our understanding of ecosystem change over decadal to millennial time-scales (Brown et al., 2023).

### Conclusion

By combining palynological diversity, REVEALS model and functional trait analysis, we disclosed detailed vegetation changes from Erhai region, southwestern China, during the climate transition period of MIS3 to LGM. Our results manifested that the Erhai region underwent a vegetation transition from temperate deciduous broadleaf forest dominance in late MIS3 to cold coniferous and mixed broadleaved/coniferous forests during the LGM, consistent with regional palynological records across the southwestern China. This vegetation dynamic involved functional compositions shifts. The competitive-driven functional convergence to partial recovery via niche differentiation during the late MIS3 were disclosed, with a gradual transition toward resource-conservative strategies responding to climate colling progress. In contrast, the vegetation transformed into a low-diversity but functional differentiation state through trait complementarity and diversification strategies during the cold LGM. Our results likely support a function-mediated climate filtering process whereby climate change regulated long-term vegetation dynamics during the MIS3–LGM transition primarily through shifts in CWM functional composition. These findings highlight the importance of incorporating community functional dynamics into studies of vegetation change on varied time scales.

## CRediT author statement

K.L.: Conceptualization, Investigation, Formal analysis, Writing - Original Draft, Writing - Review & Editing.

Z.H., P.L., X.Z., L.L.: Data curation, Formal analysis, Writing - Review & Editing. Z.T., Y.W.: Writing - Review & Editing.

M.L., J.N.: Conceptualization, Supervision, Resources, Formal analysis, Writing - Review & Editing.

## Acknowledgement

This work is financially supported by National Natural Science Foundation of China (Grant No. 42477484), and State Key Laboratory for Vegetation Structure, Function and Construction (VegLab, Grant No. VegLabOF2025006).

## Supplementary materials

Kai Li & Mengna Liao

### Regional settings

Erhai Lake (located at 99°32’-100°27’E, 25°25’-26°16’N, at an elevation of 1974 m above sea level) is a freshwater lake located in the Dali Bai Autonomous Prefecture, Yunnan Province, Southwest China (Fig. 1). The lake has a surface area of 249 km^2^ and a catchment area of 2565 km^2^. The average depth of Erhai Lake is 10.2 m, and the volume is 25.31 × 10^8^ m^3^ (Wang and Dou, 1998). The lake water is mainly supplied by precipitation (between May and October) and runoffs, and it drains southwestwards via the Xier River into the Lancang River (Me-kong) (Wang and Dou, 1998). The study region is characterized by a subtropical plateau monsoon climate, with mean annual temperature (MAT) and mean annual precipitation (MAP) of 18.0°C and 928 mm, respectively.

Dominant zonal vegetation on the Yunnan Plateau primarily comprises evergreen broadleaved forests (EBLF) and Yunnan Pine forests, which are distributed at elevations of 1000–2400 m a.s.l. and dominated by *Fagaceae* and *Pinus*. From 2400 to 2700 m a.s.l., the vegetation types give way to conifers and deciduous broadleaved trees. *Tsuga* forest (TSUF) appeared at 2300–3200 m a.s.l., as pure forest or generally mixed with temperate deciduous broadleaved forest (DBLF) (Tang, 2015). Additionally, evergreen sclerophyll *Quercus* forest (ESQF) are common and/or dominant in a belt centered between 2500 and 3600 m a.s.l. in the southeastern parts of the Tibetan Plateau and SW China (Yang et al., 2009). Cold mixed forests (CLMX), primarily comprising *Abies*, *Picea*, and *Pinus*, are predominantly distributed between 3300 and 4200 m a.s.l. As elevation continues to increase, alpine shrubland, meadow and tundra (ALSM) are prevalent, characterized by dominant species from the Cyperaceae, Poaceae, and Ranunculaceae.

**Figure S1.**
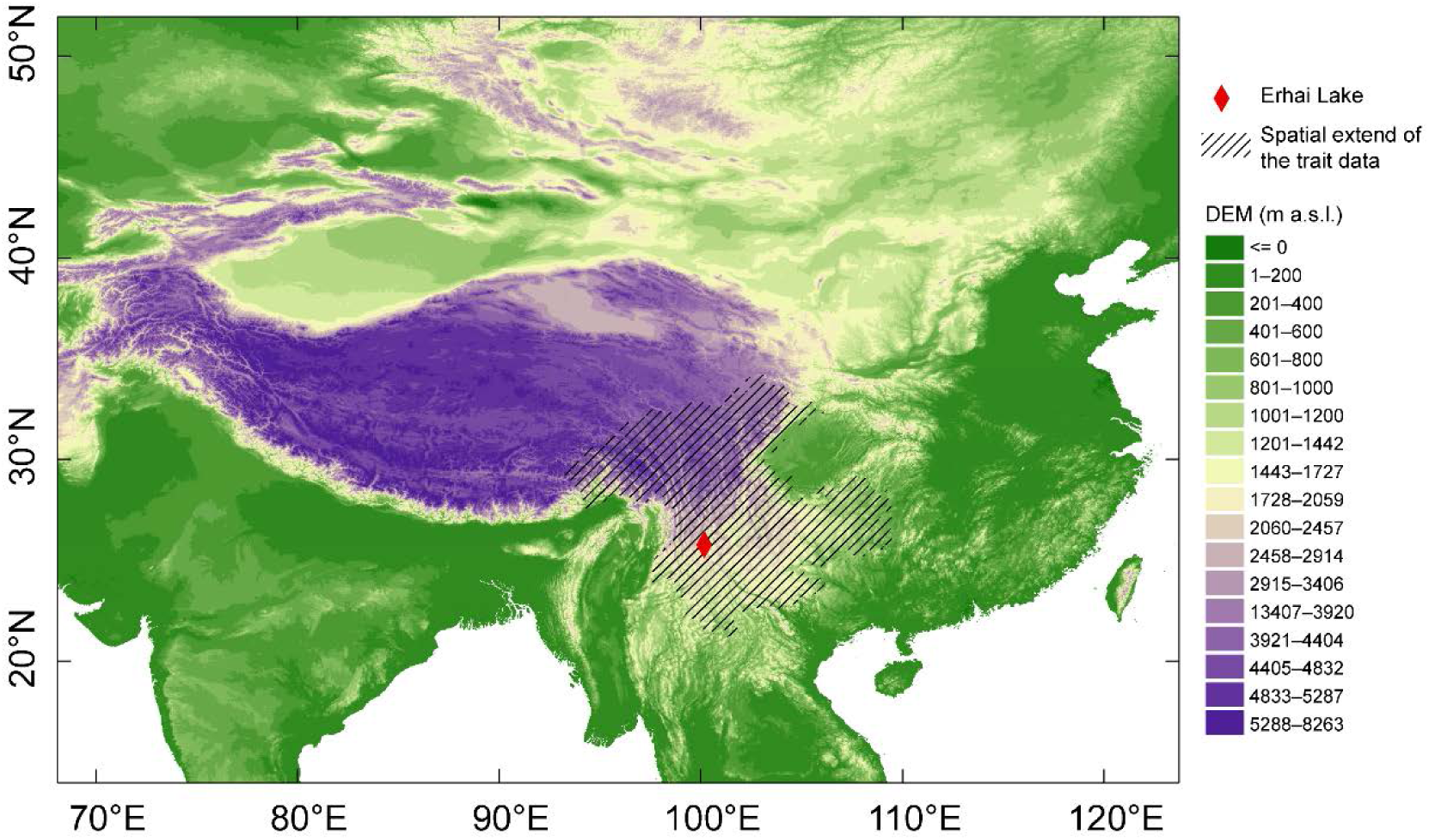
Distribution of plant trait data of modern vegetation.

**Figure S2.**
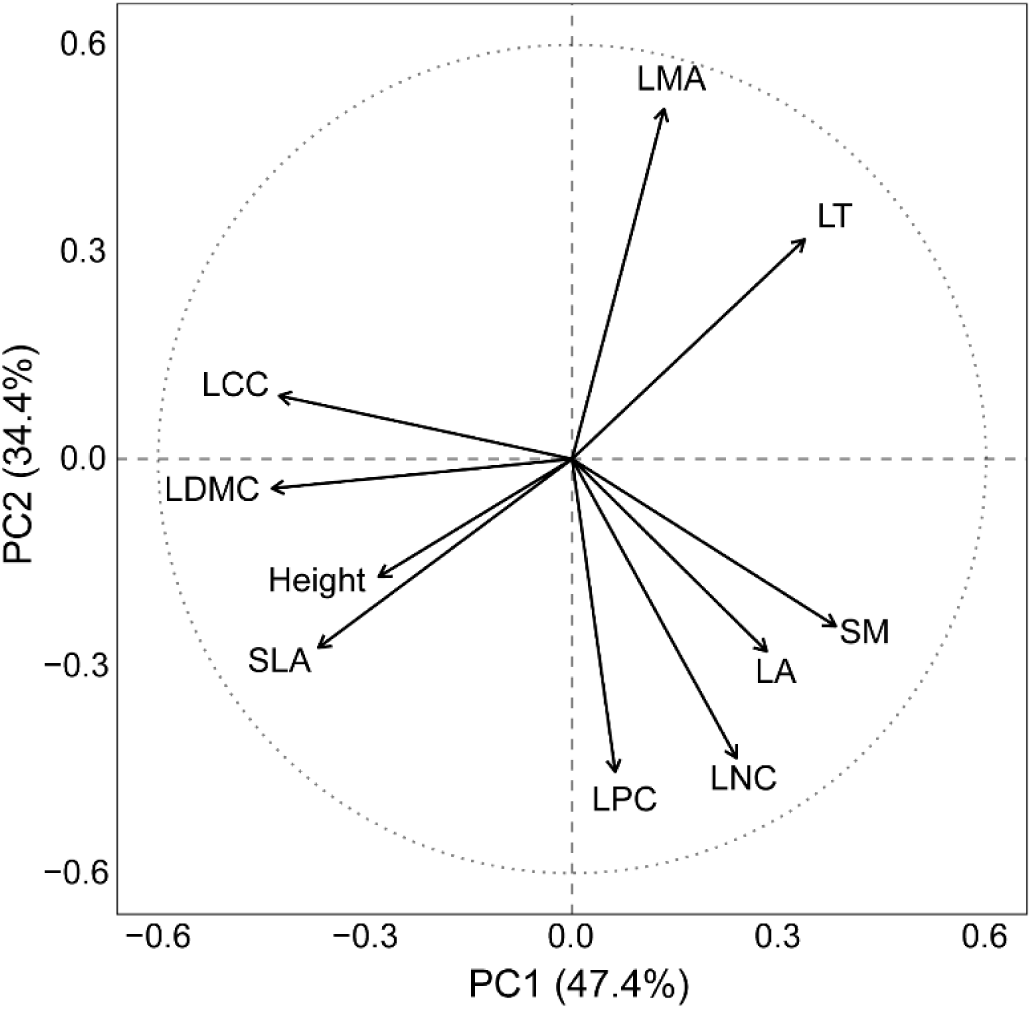
PCA of CWM functional composition.

**Figure S3.**
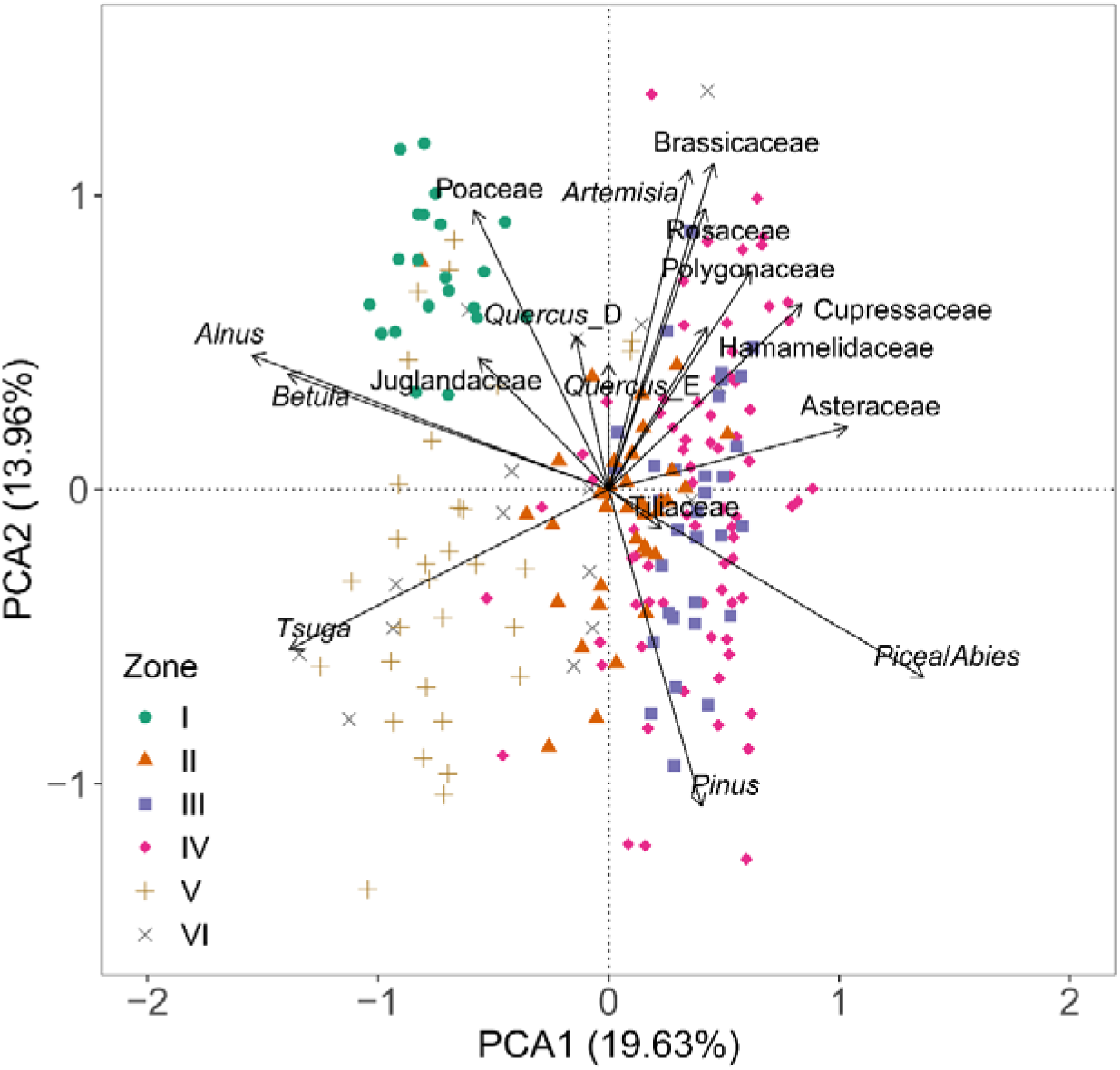
PCA of plant coverage.

**Table S1.** The Relative pollen productivity (RPP), standard errors (SE), and fall speed values of 34 selected taxa in this study.

| Pollen Taxa | RPP | SE | Long/short<br>axis<br>($\mu\text{m}$) | | Fall speed<br>(m/s) | References for RPP |
| --- | --- | --- | --- | --- | --- | --- |
| *Alnus* | 9.86 | 0.090 | 30 | 30 | 0.027 | Zhang et al., 2021 |
| Apiaceae | 12.00 | 0.250 | 35 | 15 | 0.016 | Ma et al., 2025 |
| *Artemisia* | 26.09 | 2.036 | 34 | 33 | 0.034 | Ma et al., 2025 |
| Asteraceae | 3.93 | 0.261 | 30 | 30 | 0.027 | Ma et al., 2025 |
| *Betula* | 10.26 | 0.120 | 28 | 26 | 0.021 | Zhang et al., 2021 |
| Brassicaceae | 4.60 | 0.317 | 26 | 25 | 0.019 | Ma et al., 2025 |
| *Cannabis* | 16.43 | 1.000 | 25 | 25 | 0.019 | Li et al., 2020 |
| *Castanea* | 11.49 | 0.490 | 22 | 15 | 0.010 | Li et al., 2020 |
| *Castanopsis-*<br>*Lithocarpus* | 15.47 | 0.330 | 25 | 19 | 0.014 | Zhang et al., 2021 |
| Chenopodiaceae | 7.86 | 1.998 | 30 | 30 | 0.027 | Ma et al., 2025 |
| Convolvulaceae | 0.18 | 0.030 | 70 | 70 | 0.148 | Li et al., 2020 |
| Cupressaceae | 1.11 | 0.090 | 40 | 40 | 0.048 | Li et al., 2020 |
| Cyperaceae | 10.72 | 0.021 | 53 | 38 | 0.061 | Ma et al., 2025 |
| *Ephedra* | 3.30 | 0.230 | 50 | 25 | 0.038 | Ma et al., 2025 |
| Fabaceae | 1.67 | 0.096 | 39 | 38 | 0.044 | Ma et al., 2025 |
| Gentianaceae | 6.66 | 0.416 | 38 | 33 | 0.038 | Ma et al., 2025 |
| Hamamelidaceae | 2.95 | 0.022 | 37 | 36 | 0.040 | Chen et al., 2019 |
| *Humulus* | 16.43 | 1.000 | 30 | 30 | 0.027 | Li et al., 2020 |
| Juglandaceae | 4.83 | 0.180 | 39 | 36 | 0.041 | Zhang et al., 2021 |
| Labiatae | 1.24 | 0.190 | 38 | 36 | 0.041 | Li et al., 2020 |
| Liliaceae | 1.49 | 0.110 | 42 | 25 | 0.032 | Li et al., 2020 |
| Oleaceae | 3.94 | 0.730 | 39 | 25 | 0.029 | Li et al., 2020 |
| *Picea/Abies* | 29.40 | 0.870 | 52 | 50 | 0.077 | Zhang et al., 2021 |
| *Pinus* | 16.68 | 0.510 | 37 | 36 | 0.040 | Wieczorek & Herzsuh, 2020 |
| *Plantago* | 48.75 | 1.000 | 30 | 30 | 0.027 | Ma et al., 2025 |
| Poaceae | 1.00 | 0.000 | 23 | 23 | 0.016 | Ma et al., 2025 |
| Polygonaceae | 8.06 | 0.346 | 48 | 35 | 0.051 | Ma et al., 2025 |
| *Quercus_D* | 4.27 | 0.040 | 35 | 31 | 0.031 | Zhang et al., 2021 |
| *Quercus_E* | 5.19 | 0.070 | 28 | 28 | 0.024 | Zhang et al., 2021 |
| Rosaceae | 0.67 | 0.012 | 30 | 24 | 0.022 | Chen et al., 2019; Guo, 2021; Jiang<br>et al., 2020 |
| Rubiaceae | 1.23 | 0.360 | 26 | 25 | 0.019 | Li et al., 2020 |
| Tiliaceae | 0.65 | 0.110 | 40 | 40 | 0.048 | Li et al., 2020 |
| *Tsuga* | 22.31 | 3.200 | 44 | 44 | 0.058 | Chaput and Gajewski, 2018 |
| *Ulmus* | 4.13 | 0.920 | 34 | 34 | 0.035 | Li et al., 2020 |

**Table S2.**
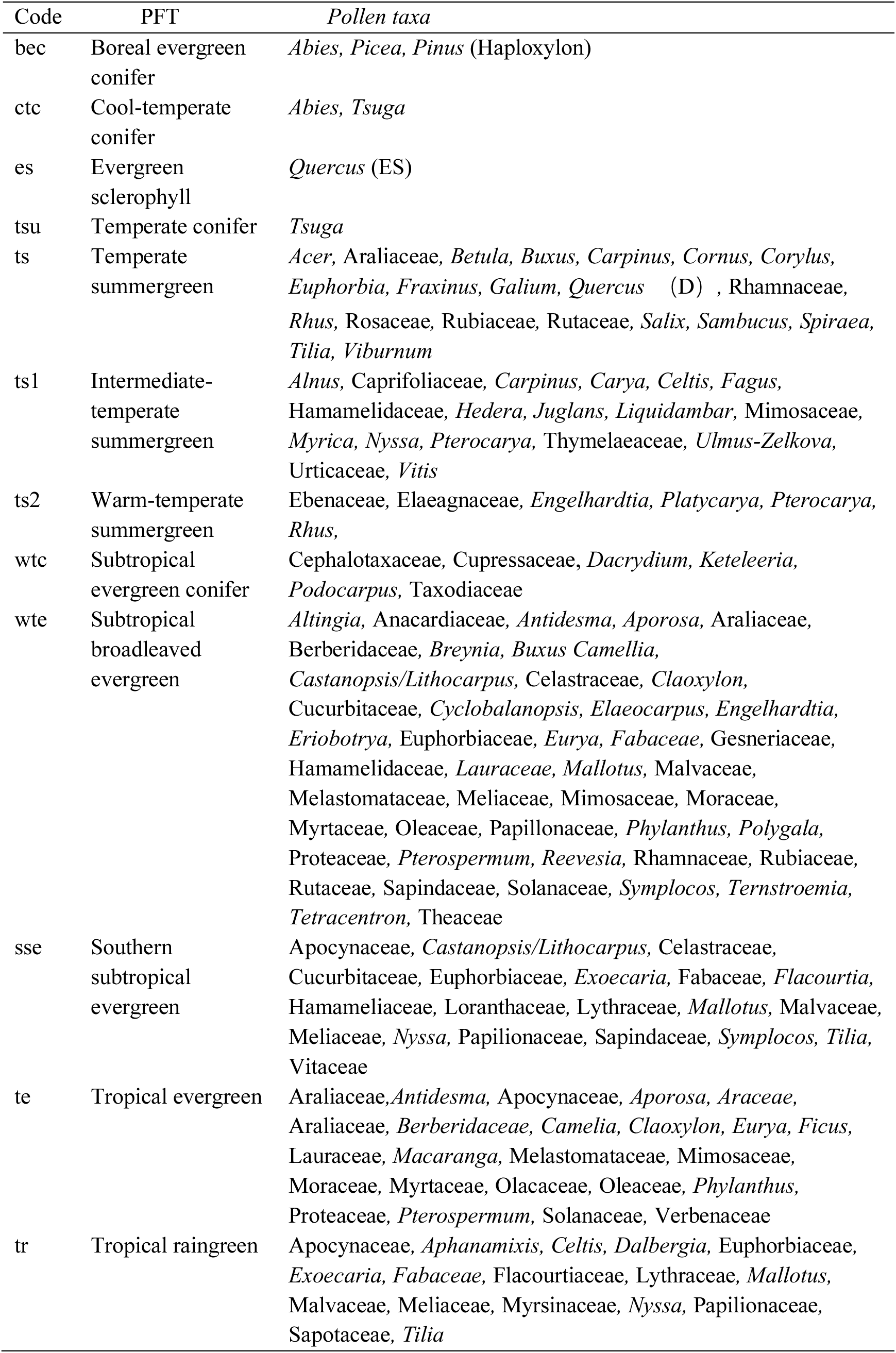
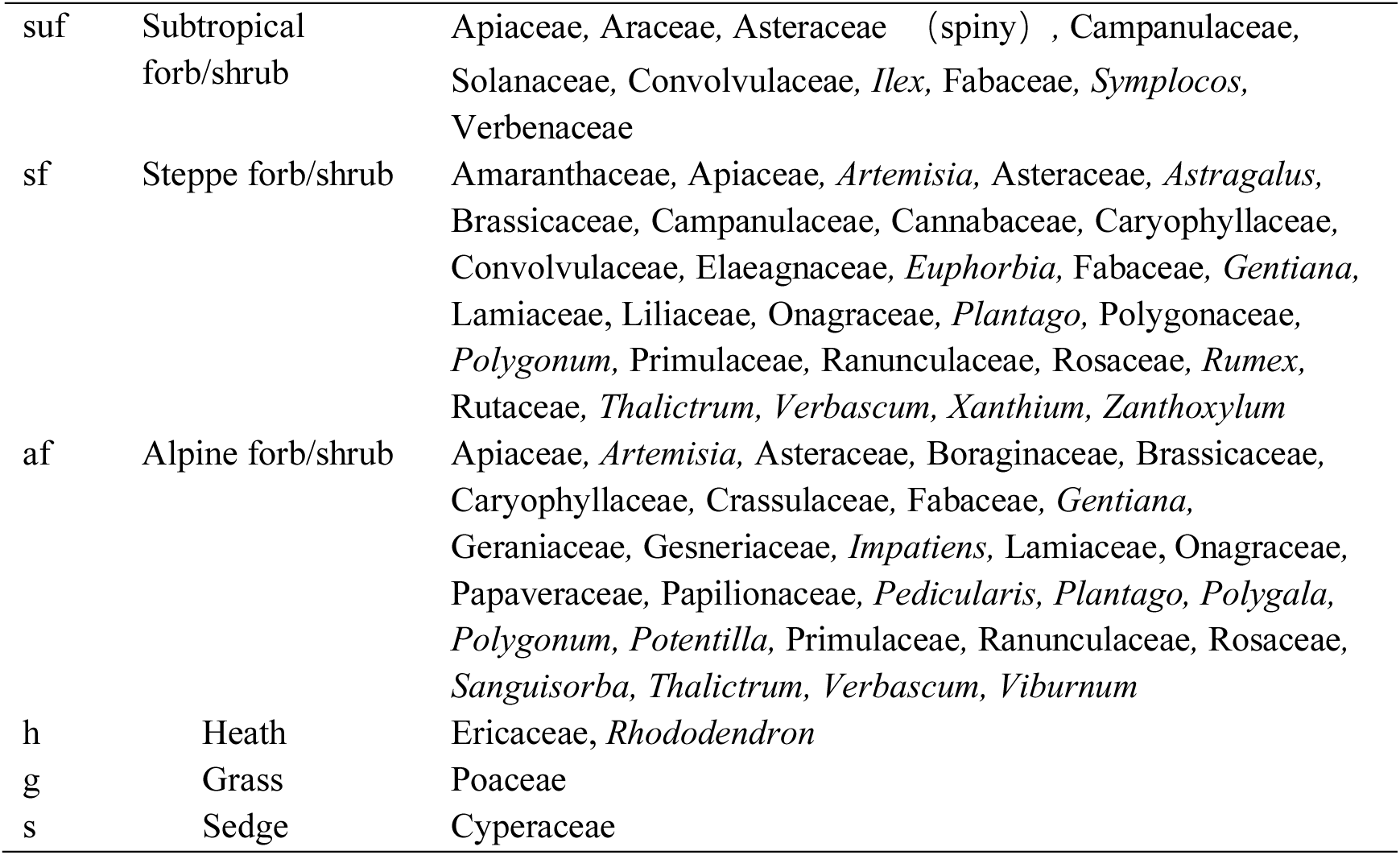
Taxa to Plant function types (PFTs), modified from Ni et al. (2014) and Zheng et al. (2023).

| Code | PFT | Pollen taxa |
| --- | --- | --- |
| bec | Boreal evergreen<br>conifer | *Abies, Picea, Pinus* (Haploxyton) |
| ctc | Cool-temperate<br>conifer | *Abies, Tsuga* |
| es | Evergreen<br>sclerophyll | *Quercus* (ES) |
| tsu | Temperate conifer | *Tsuga* |
| ts | Temperate<br>summergreen | *Acer, Araliaceae, Betula, Buxus, Carpinus, Cornus, Corylus, Euphorbia, Fraxinus, Galium, Quercus* (D) , *Rhamnaceae, Rhus, Rosaceae, Rubiaceae, Rutaceae, Salix, Sambucus, Spiraea, Tilia, Viburnum* |
| ts1 | Intermediate-<br>temperate<br>summergreen | *Alnus, Caprifoliaceae, Carpinus, Carya, Celtis, Fagus, Hamamelidaceae, Hedera, Juglans, Liquidambar, Mimosaceae, Myrica, Nyssa, Pterocarya, Thymelaeaceae, Ulmus-Zelkova, Urticaceae, Vitis* |
| ts2 | Warm-temperate<br>summergreen | *Ebenaceae, Elaeagnaceae, Engelhardtia, Platycarya, Pterocarya, Rhus,* |
| wtc | Subtropical<br>evergreen conifer | *Cephalotaxaceae, Cupressaceae, Dacrydium, Keteleeria, Podocarpus, Taxodiaceae* |
| wte | Subtropical<br>broadleaved<br>evergreen | *Altingia, Anacardiaceae, Antidesma, Aporosa, Araliaceae, Berberidaceae, Breynia, Buxus Camellia, Castanopsis/Lithocarpus, Celastraceae, Claoxylon, Cucurbitaceae, Cyclobalanopsis, Elaeocarpus, Engelhardtia, Eriobotrya, Euphorbiaceae, Eurya, Fabaceae, Gesneriaceae, Hamamelidaceae, Lauraceae, Mallotus, Malvaceae, Melastomataceae, Meliaceae, Mimosaceae, Moraceae, Myrtaceae, Oleaceae, Papillonaceae, Phyllanthus, Polygala, Proteaceae, Pterospermum, Reevesia, Rhamnaceae, Rubiaceae, Rutaceae, Sapindaceae, Solanaceae, Symplocos, Ternstroemia, Tetracentron, Theaceae* |
| sse | Southern<br>subtropical<br>evergreen | *Apocynaceae, Castanopsis/Lithocarpus, Celastraceae, Cucurbitaceae, Euphorbiaceae, Exoecaria, Fabaceae, Flacourtia, Hamameliaceae, Loranthaceae, Lythraceae, Mallotus, Malvaceae, Meliaceae, Nyssa, Papilionaceae, Sapindaceae, Symplocos, Tilia, Vitaceae* |
| te | Tropical evergreen | *Araliaceae, Antidesma, Apocynaceae, Aporosa, Araceae, Araliaceae, Berberidaceae, Camellia, Claoxylon, Eurya, Ficus, Lauraceae, Macaranga, Melastomataceae, Mimosaceae, Moraceae, Myrtaceae, Olacaceae, Oleaceae, Phyllanthus, Proteaceae, Pterospermum, Solanaceae, Verbenaceae* |
| tr | Tropical raingreen | *Apocynaceae, Aphanamixis, Celtis, Dalbergia, Euphorbiaceae, Exoecaria, Fabaceae, Flacourtiaceae, Lythraceae, Mallotus, Malvaceae, Meliaceae, Myrsinaceae, Nyssa, Papilionaceae, Sapotaceae, Tilia* |
| suf | Subtropical forb/shrub | Apiaceae, Araceae, Asteraceae (spiny) , Campanulaceae, Solanaceae, Convolvulaceae, *Ilex*, Fabaceae, *Symplocos*, Verbenaceae |
| sf | Steppe forb/shrub | Amaranthaceae, Apiaceae, *Artemisia*, Asteraceae, *Astragalus*, Brassicaceae, Campanulaceae, Cannabaceae, Caryophyllaceae, Convolvulaceae, Elaeagnaceae, *Euphorbia*, Fabaceae, *Gentiana*, Lamiaceae, Liliaceae, Onagraceae, *Plantago*, Polygonaceae, *Polygonum*, Primulaceae, Ranunculaceae, Rosaceae, *Rumex*, Rutaceae, *Thalictrum*, *Verbascum*, *Xanthium*, *Zanthoxylum* |
| af | Alpine forb/shrub | Apiaceae, *Artemisia*, Asteraceae, Boraginaceae, Brassicaceae, Caryophyllaceae, Crassulaceae, Fabaceae, *Gentiana*, Geraniaceae, Gesneriaceae, *Impatiens*, Lamiaceae, Onagraceae, Papaveraceae, Papilionaceae, *Pedicularis*, *Plantago*, *Polygala*, *Polygonum*, *Potentilla*, Primulaceae, Ranunculaceae, Rosaceae, *Sanguisorba*, *Thalictrum*, *Verbascum*, *Viburnum* |
| h | Heath | Ericaceae, *Rhododendron* |
| g | Grass | Poaceae |
| s | Sedge | Cyperaceae |

**Table S3.** Assignments of plant functional types to biomes, modified from Ni et al. (2014) and Zheng et al. (2023).

| Code | Biome | Plant functional type (PFT) |
| --- | --- | --- |
| ALSM | Alpine shrub-land, meadow/tundra | af, g, h, s |
| CLMX | Cold mixed forest | bec, ctc |
| ESQF | Evergreen sclerophyll forest | es |
| TSUF | *Tsuga* forest | tsu |
| DBLF | Temperate deciduous broadleaved forest | h, ts, ts1, ts2 |
| EBLF | Subtropical evergreen broadleaved forest | te, tr, ts2, wtc, wte |
| TRFO | Tropical rainforest | te, sse, suf |

**Table S4.** The plant functional traits used in this study and their ecological significance.

| Plant function traits<br>& References | Changes and drivers |
| --- | --- |
| Plant height<br>(Falster and Westoby 2003) | The height of a community is directly related to the vegetation composition of the community. Communities dominated by trees generally have taller vegetation than those dominated by herbs. Deciduous trees have taller heights as compared to evergreen trees. Taller trees can occupy favorable positions, receive more sunlight and carbon dioxide, and consequently, photo-synthesize more, grow faster, and form taller canopies. |
| Seed mass (SM)<br>(Saatkamp et al., 2019) | Seed mass is intimately linked to reproductive strategy, serving as a key index of variation in maternal investment. Plants must navigate a fundamental trade-off between seed size and number, giving rise to a continuum of strategies ranging from few large seeds to many small ones. Moreover, large seeds generally withstand harsh environmental conditions more effectively than their smaller counterparts. |
| Leaf thickness<br>(LT)<br>(Nobel and Hartsock 1981, Gratani et al. 2018) | Thicker leaves can impede light penetration, thereby hampering its reach to the deeper cells of the leaf, leading to a decline in the photosynthetic capacity of those cells. Nonetheless, a greater leaf thickness may also curb water transpiration, thereby prompting plants to use and store water more effectively in water-scarce environments. Besides, thicker leaves can lower the damage caused by adversity factors and scale up the plant's resistance towards unfavorable conditions. Thicker leaves can typically store more nutrients and organic matter, enabling plants to maintain regular growth even in conditions of nutrient deficiency or severe environmental changes. |
| Leaf area<br>(LA)<br>(Fang et al. 2019) | A greater LA facilitates increased light energy absorption, photosynthesis efficiency, water uptake capacity, and nutrient acquisition. Consequently, plant growth rate, biomass accumulation, and nutrient utilization efficiency improve. Conversely, smaller leaf area decreases transpiration and water loss whilst improving water use efficiency and facilitating adaptation to water-stressed environments. |
| Specific leaf area<br>(SLA)<br>(Wilson et al. 1999) | Plants with higher SLA require more sunshine and nutrients, indicating they are better suited to environments with sufficient resources. Conversely, plants with lower SLAs have lower requirements for light and nutrients, indicating they are more adapted to environments with limited resources. |
| Leaf mass per area<br>(LMA)<br>(Poorter et al., 2009; Villar et al., 2013) | Higher LMA reflects an increase in the construction cost per unit leaf area. Plants enhance their tolerance in cold and nutrient-constrained environments by forming thicker, denser, and more mechanically strong leaves. |
| Leaf dry matter content<br>(LDMC)<br>(Wilson et al. 1999) | Plants with higher LDMC are generally considered to have higher leaf structural strength and tolerance to adversity, adapting to nutrient-poor and drought conditions. Plants with lower LDMC, on the other hand, usually have higher photosynthetic efficiency and are adapted to adequate light conditions. |
| Leaf carbon content<br>(LCC)<br>(Prieto et al. 2018) | Higher LCC is usually associated with a resource-conservative growth strategy, meaning that plants tend to undergo more efficient carbon fixation with limited resource availability in order to reduce resource consumption. This growth strategy is usually characterized by higher LDMC and lower SLA, as well as lower leaf N, P and K concentrations. |
| Leaf nitrogen content<br>(LNC)<br>(Kattge et al. 2009) | Plants with a high LNC typically exhibit enhanced photosynthetic efficiency and growth rates, which is a successful strategy in nutrient-rich environments. This increase in photosynthetic enzyme activity and carbon fixation capacity in leaves is due to the high amount of leaf nitrogen, resulting in an improved light energy use efficiency and biomass accumulation. Such a growth strategy is commonly employed by plants that grow fast and compete favorably, including fast-growing annuals and pioneer species. |
| Leaf phosphorus<br>content (LPC)<br>(Gong et al. 2020);<br>Reich and Oleksyn.,<br>2004) | LPC is positively correlated with mean annual precipitation and mean annual temperature; High LPC indicate that plants strengthened nutrient acquisition capacity when the climate became slightly milder, exhibiting a brief resource-acquisitive strategy. |

